# Deciphering the evolution of sex determination across the termite tree of life using high-quality genome assemblies

**DOI:** 10.64898/2025.12.29.696834

**Authors:** Simon Hellemans, Yi-Ming Weng, Mara J. Julseth, Kokuto Fujiwara, Cédric Aumont, Cong Liu, Alina A. Mikhailova, Aleš Buček, Esra Kaymak, David Sillam-Dussès, Yves Roisin, Jan Šobotník, Kiyoto Maekawa, Mark C. Harrison, Dino P. McMahon, Thomas Bourguignon

## Abstract

Most termites exhibit a unique sex-determining system, featuring multiple X and Y chromosomes that fuse into large complexes during male meiosis. The evolutionary origins of such complexes remain largely unknown. Using the genomes of 45 termites and two cockroaches, we investigated the evolution of sex determination systems in an entire insect lineage. We found that termite sex chromosomes are largely undifferentiated, likely reflecting extensive ongoing recombination. Evolving from the X0 system of cockroaches, most early-diverging termites exhibit a Y chromosome bearing the sex-determining gene *doublesex*, whereas *doublesex* is autosomal in most other termites. In contrast, other species exhibit multiple sex chromosomes that have undergone frequent turnover, except for Termitidae, which harbor conserved sex chromosomes. Our findings reveal important reworkings of the ancestral *transformer*-*doublesex* pathway in termites and suggest a potential role of *doublesex* in the formation of meiotic chromosomal complexes and caste differentiation.

## Introduction

Sex determination mechanisms are remarkably diverse amongst animals (*1–7*). In insects, sex is generally determined genetically, or more rarely by environmental factors (*8*, *9*). The most common sex determination system (SDS) is based on the heterogamy of one gene, a full chromosome, or a fraction of a chromosome (*9*), with the heterogametic sex being female in ZZ/ZW SDS and male in XX/XY SDS (*1*, *10*), as is the case in most termites.

Sex chromosomes are diverse and have evolved hundreds of times independently across animals (*1*). Young sex chromosomes are typically homomorphic, exhibiting limited recombination suppression, with homologues appearing morphologically similar and presenting low sequence divergence (*1*, *11*, *12*). In contrast, ancient sex chromosomes are generally heteromorphic, with extensive divergence in sequence identity, gene content, and morphology due to reduced recombination driving the degradation of the repetitive-rich Y and W chromosomes (*1*, *13–16*). Sex chromosomes are occasionally conserved between distantly related taxa, as exemplified by *Drosophila* and the German cockroach *Blattella germanica*, which have homologous X chromosomes despite 400 million years of divergence (*17*). However, various mechanisms can alter the identity of sex chromosomes, including the translocation of the sex-determining gene on an autosome, the evolution of a new sex-determining gene, or the recruitment of an autosome via chromosomal rearrangements (*1*, *18*), which leads to sex chromosome turnover across lineages.

As for sex chromosomes, the sex-determining signal and its downstream cascade of gene activation vary significantly across animals, although some components are largely conserved. In insects, the canonical sex determination cascade revolves around the *transformer*-*doublesex* (*tra*-*dsx*) axis (*1*, *19–21*). In many insects, the RNA splicing factor *tra* evolved to promote the female-specific splicing of *dsx*, which induces the development of female traits, while the presence of full-length *dsx* leads to male differentiation (*20*, *22*, *23*). The upstream activators of these genes vary greatly amongst taxa, often consisting of male- or female-specific genes or genes differentially expressed between sexes (*1*, *10*). For example, sex-lethal (*Sxl*) initiates female development by positively regulating *tra* in *Drosophila* (*15*, *24*), while the M-factor represses *tra* transcription to trigger male development in the housefly *Musca domestica* (*25*). Sex-determining genes can occur on either autosomes or sex chromosomes (*10*, *15*, *26*, *27*). For instance, the M factor is located either on the X or Y chromosomes or the autosomes in *M. domestica* (*25*). The precise molecular mechanisms of sex determination remain unknown in many insects, including termites.

Termites (Isoptera) are eusocial cockroaches (Blattodea), sister to the subsocial woodroach *Cryptocercus* (*28*). Termite genomes have several modes of expression that generate workers, soldiers, and reproductives differing in their anatomy, physiology, and behavior. Non-termite cockroaches are diploid in both sexes, except for males that are hemizygous for the sex chromosome (XX/X0 SDS). In contrast, all termites but the XX/X0 *Stolotermes* have an XX/XY SDS (*29*, *30*), with karyotypically homomorphic sex chromosomes (*31*, *32*). Interestingly, most species of Icoisoptera (Kalotermitidae + Neoisoptera) exhibit chains or rings involving multiple sex chromosomes during male meiosis (*30*, *33–36*), a rare feature also reported in platypus (*36*) and an Amazonian frog (*37*). Another unique feature of most termites is the loss of the sex-spliced OD2 domain of the sex-determining gene *dsx*, with OD1 being expressed in males only (*38–41*). How termite sex chromosomes and their determination mechanisms have evolved remains poorly understood.

Here, we resolved the evolution of termite SDS by identifying sex chromosomes and sex-determining genes across a representative set of species. We identified the sex chromosomes in 36 of the high-quality genomes of 45 termites and two cockroaches recently published by Liu *et al*. (*42*) using sex-specific short reads generated in this study. We also identified sex-determining genes in all 47 genomes. Using this dataset, we showed that termite sex chromosomes (*i*) exhibit sex-specific differentiation despite their homomorphic morphology; (*ii*) do not carry the genes of the canonical sex-determining cascade, except in a few early diverging termite lineages, and (*iii*) experienced turnover across lineages, except in Termitidae, in which they are fixed. Finally, (*iv*) we explored the evolutionary trajectory of *dsx* and the genes it regulates.

## Results and Discussion

### Termite genomes exhibit minimal sexual differentiation, with largely undifferentiated sex chromosomes

We generated sex-specific short reads at a sequencing depth of at least 15x for 34 termites and two cockroaches (*Blatta orientalis* and *Cryptocercus meridianus*) with a known genome (Supplementary Data 1). We identified sex chromosomes using six computational methods relying on comparative male-female coverage (*C_MF_*), single nucleotide polymorphism (SNPs), and k-mers (Supplementary Data 1-4; Dryad: Files 1-2). We found that about 6% of both cockroach genomes was sex-linked (Figure 1F; Supplementary Data 4; Supplementary Figures 1,2; Dryad: File 2). Sex-linked scaffolds exhibited higher coverage in females than in males (negative log_2_(*C_MF_*)), without any within-scaffold shifts in differential coverage (Supplementary Figure 3) nor male-specific SNPs (Supplementary Figure 4). This pattern is consistent with an XX/X0 SDS (Figures 1, 2 A-C), with hemizygous males and diploid females for the X chromosome (*17*, *29*). In termites, between 0.02% and 34.15% of the genome was identified as sex-linked (Figure 1F; Supplementary Data 4; Supplementary Figure 1). Termite scaffolds generally exhibited similar coverages in both males and females (Supplementary Figures 1,2; Dryad: File 2), indicating that termite X and Y chromosomes present limited differentiation and co-assemble. As all termites have sex chromosomes, the very low proportion of sex-linked scaffolds in some termite genomes reflects our inability to identify homomorphic sex chromosomes in these species rather than an actual absence of sex chromosomes.

**Figure 1:**
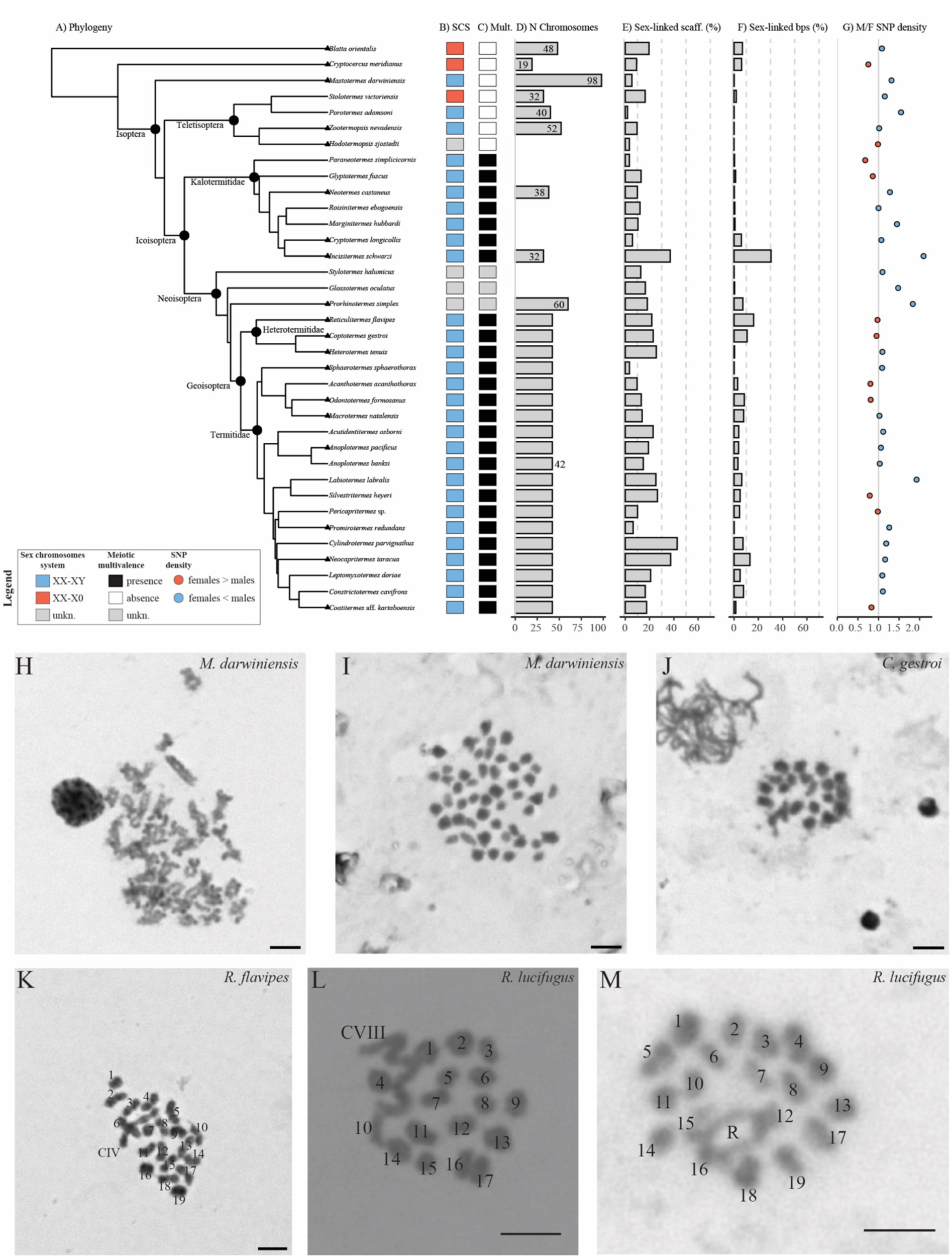
Sex chromosomes in Isoptera. (**A**) Phylogenetic relationships among the 39 species for which sex-specific reads were obtained (tree simplified from Liu *et al*. (*42*)). Species with chromosome-level assemblies are marked by a triangle (Supplementary Data 1). Sex determination systems (SDS) of studied species: (**B**) systems of sex chromosomes (XX/XY, blue; XX/X0, red; unknown, grey), (**C**) presence/absence of a multivalent chain or ring during metaphase I in males (presence, black; absence, white; unknown, grey), and (**D**) number of chromosomes. Percentages of consensus sex-linked (**E**) scaffolds and (**F**) base pairs. (**G**) Male-to-female SNP density deviation from the median number of SNPs weighted by scaffold sizes (higher SNP density in males, blue; higher SNP density in females, red). (**H**–**M**) Spreads of chromosomes in meiosis I in males (scalebars, 5 μm). Pictures of *Mastotermes darwiniensis* chromosomes showing (**H**) multiple chiasmata on all chromosomes indicative of recombination and (**I**) the 49 bivalents (2*n* = 98). (**J**) Picture of the 21 bivalents of *Coptotermes gestroi*, including a V-shaped bivalent. (**K**) Picture of the 19 bivalents and a multivalent chain CIV of *Reticulitermes flavipes*. (**L**, **M**) Pictures of the chromosomes of *Reticulitermes lucifugus corsicus* exhibiting either (**L**) 17 bivalents and a multivalent chain CVIII or (**M**) 19 bivalents and a multivalent ring RIV.

**Figure 2:**
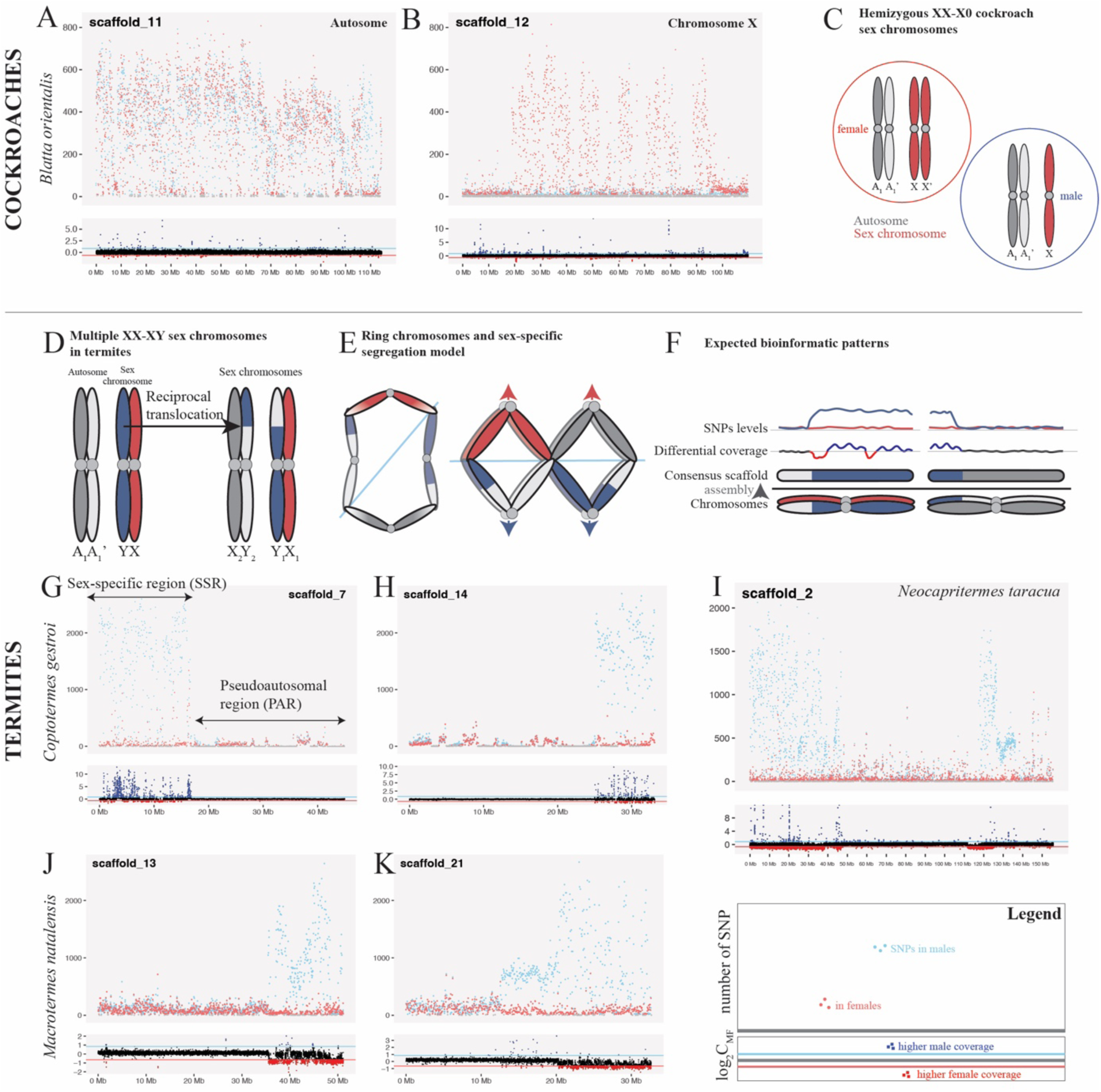
Sex-specific chromosomal patterns. (**A**–**B**, **G**–**I**) Sliding SNP counts (top; females, red; males, blue) and male-to-female log_2_(*C_MF_*) coverage (bottom; higher coverage in females, red; in males, blue) along the length of selected scaffolds. (**A**) An autosome and (**B**) the X chromosome in the XX/X0 cockroach *B. orientalis*. The autosome exhibits similar levels of SNPs in males and females, while the X chromosome exhibits elevated SNPs in females only. (**C–F**) Schematic diagrams illustrating (**C**) cockroach XX/X0 sex chromosomal system with males hemizygous only for the X chromosomes and (**D–F**) multiple XX/XY SDS in termites. (**D**, **E**) Reciprocal translocation between the canonical sex chromosome and an autosome explaining the formation of a quadrivalent ring and sex-specific inheritance of X and Y chromosomes during the male meiosis, following the models of Vincke & Tilquin (*33*) and Pajpach *et al*. (*59*). (**F**) Expected bioinformatic patterns under the translocation model, with two of the top 21 scaffolds exhibiting elevated SNP counts in males (blue) and male-to-female coverage diverging from equality. (**G**–**I**) Examples of sex chromosomes in Geoisoptera, in (**G**, **H**) the Heterotermitidae *Coptotermes gestroi* and the Termitidae (**J**, **K**) *Macrotermes natalensis* and (**I**) *Neocapritermes taracua*. Chromosome portions with elevated male-to-female coverage and SNPs in males are, non-recombining, sex-specific arms (strata), while portions with equal coverage and low SNP counts recombine between sexes. Elevated male-to-female coverage in (**G**, **H**) highlights that the scaffold included in the reference assembly is closer in similarity to the Y-specific arm, while higher female coverage in (**J**, **K**) indicates that the reference scaffold has a higher similarity to the X-specific arm.

We further assessed the level of differentiation of the sex-linked genomic fraction in termites using dN/dS analyses. Genes located in regions of sex chromosomes where recombination is suppressed, at least partially, or under strong sex-specific selection are expected to exhibit elevated non-synonymous to synonymous substitution rates (dN/dS) (*43–45*). We compared the dN/dS values of genes located on sex chromosomes to those of autosomal genes for each species having more than 20 sex-linked genes. dN/dS values were computed for the 17,652 hierarchical orthogroups (HOGs) identified by Liu *et al*. (*42*) (Dryad: File 3). We performed two-sided *t*-tests for 28 species, 12 of which were significant, all with genes present on sex chromosomes showing higher dN/dS values than autosomal genes (Supplementary Data 6). Therefore, while some termite species present a mild differentiation of sex chromosomes, many species present no chromosome-scale genetic differentiation between sexes, which is in line with their lack of morphological differentiation in karyotypes (*29–32*, *46*). The low level of genetic differentiation between X and Y chromosomes may be explained by ongoing recombination between the X and Y chromosomes or a recent origin of sex chromosomes in many species.

### The sex chromosomes of most Kalotermitidae and Neoisoptera exhibit evolutionary strata

Our scaffold-wide analyses only identified a small fraction of genomes as sex-linked in some termite species (Figure 1), presumably because of pervasive recombination between large fractions of the X and Y chromosomes. However, some parts of sex chromosomes may have lost their ability to recombine, forming evolutionary strata —chromosomal regions where recombination between X and Y chromosomes has ceased, a hallmark of sex chromosome evolution (*12*). We searched 16 near chromosome-level genome assemblies of Icoisoptera, including three Kalotermitidae and 13 Neoisoptera (Figure 1; Supplementary Table 1), for the presence of evolutionary strata in the sex chromosomes. Twelve of these 16 genomes exhibited at least one large contig (*i.e.*, an entire sex chromosome) composed of a pseudo-autosomal region (PAR) and a sex-specific region (SSR) representing an evolutionary stratum. The SSR was easily distinguishable from the PAR by an unbalanced log_2_(*C_MF_*) and a higher SNP count in males (Figure 2G-I, Supplementary Figures 2–4). For instance, SSRs were observed in two of the 21 major scaffolds of the heterotermitid *Coptotermes gestroi* and the termitid *Macrotermes natalensis* (Figure 2G-I), which is in line with karyotypic observations of the latter two species. The karyotypes of *Coptotermes* and Termitidae (*29*) included two sex chromosomes forming a meiotic quadrivalent ring during male meiosis (Figures 1–2) (*29*, *31*). This meiotic feature likely arose from a heterologous translocation between the ancestral Y chromosome and an autosome (*33*), which generated non-recombining SSRs on two chromosome pairs (Figure 2D–F), as we found in most species of Icoisoptera (Figure 2G–K; Supplementary Figures 2–4). SSRs were observed at both ends of the longest scaffold of the termitid *Neocapritermes taracua* (Figure 2I). Since this scaffold appeared to combine three of the 21 major scaffolds (Figure 4), it possibly resulted from a mis-assembly or reflected a unique chromosome fusion in Neocapritermitinae. Overall, our results show that the sex chromosomes of Icoisoptera present evolutionary strata despite co-assembling and presenting limited chromosome-scale genetic divergence.

### The early-branching Mastotermitidae, Archotermopsidae, and Hodotermopsidae have undifferentiated sex chromosomes with male-restricted sex-determining gene dsx

Our results indicated that only a small fraction of the near chromosome-level genome assemblies of *Mastotermes darwiniensis* (Mastotermitidae), *Hodotermopsis sjosdedti* (Hodotermopsidae), and *Zootermopsis nevadensis* (Archotermopsidae) was sex-linked, with none of the assembled chromosomes exhibiting significant sex-linkage (Figure 1F; Supplementary Figures 2–4). Considering that these termites do not exhibit meiotic complexes like those found in Icoisoptera (*32*), their sex chromosomes are likely a single homomorphic autosome-looking pair. To identify their sex chromosomes, we determined the genomic location of the *pros*-*dsx*-*UXT* syntenic block, which contains the sex-determining gene *dsx* (*38*, *41*). We reasoned that the block presence on a scaffold might constitute the sole sex-specific feature in these termites. We performed our analysis on the 47 genomes of Liu *et al*. (*42*). We successfully identified *pros* and *UXT* genes in all 47 genomes and the OD1 domain of *dsx* in all genomes but those of *H. sjostedti* and *Z. nevadensis* (Supplementary Data 10; Figure 3). The *pros-dsx-UXT* synteny was conserved in the two cockroaches and 35 termite species, but it was not in 10 termite species, including *M. darwiniensis*, *H. sjostedti*, and *Z. nevadensis* (Figure 3), as previously reported for the latter (*41*, *47*), indicating a translocation of *dsx*.

**Figure 3:**
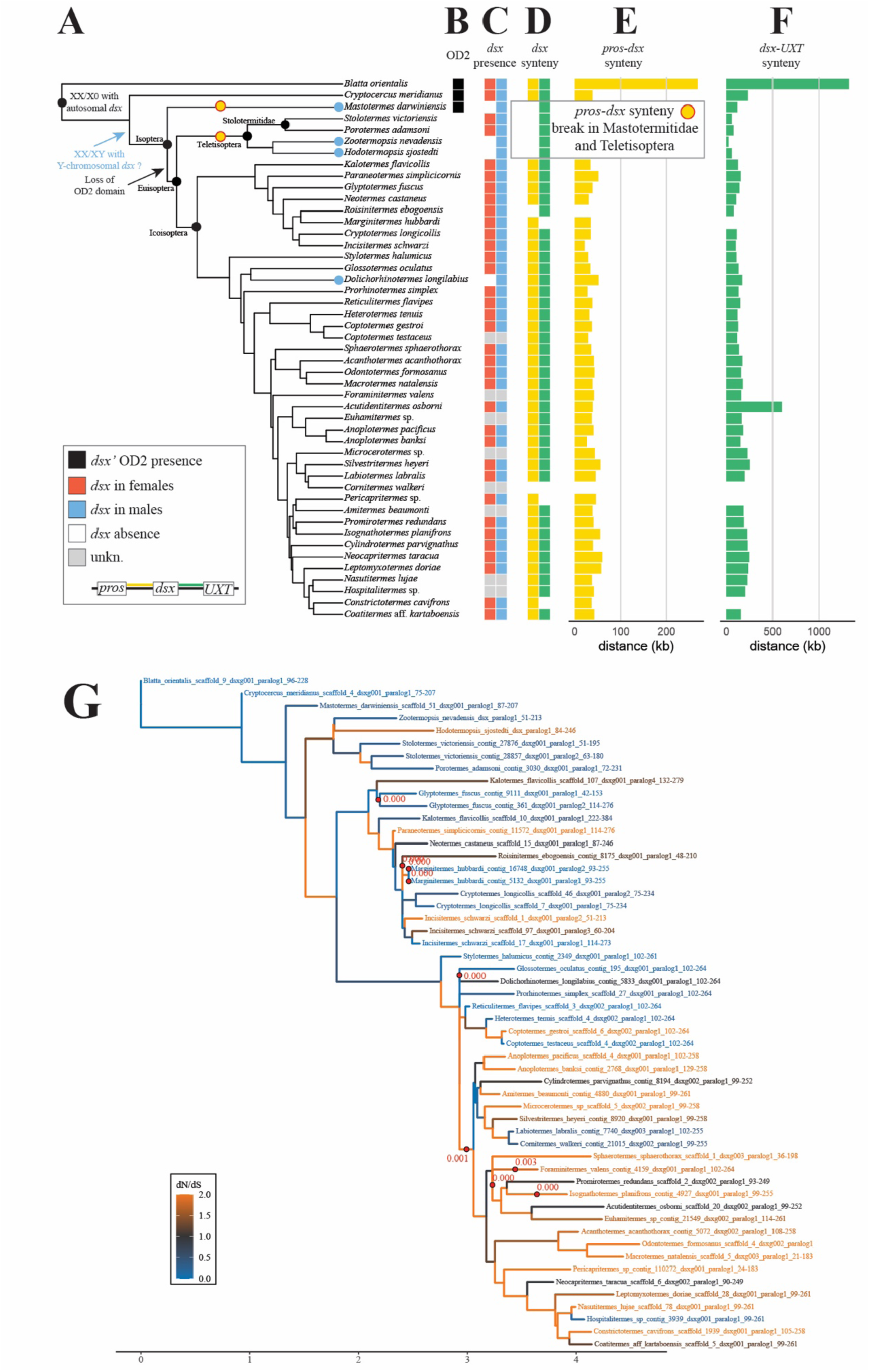
The *doublesex* gene in Isoptera. (**A**) Phylogenetic relationships among the 47 species in the genomic dataset (tree from Liu *et al*. (*42*)), annotated with *dsx* patterns. (**B**) Presence/absence of *dsx* OD2 domain in Blattodea (black, presence; white, absence). (**C**) Presence/absence of *dsx* OD1 domain in males and females (presence in females, red; males, blue; absence, white; unknown, grey). Genomic location of *doublesex* through (**D**) the *prospero*-*doublesex* (presence, gold; absence, white) and *prospero*-*UXT* (presence, green; absence, white) syntenies, and distances in k-bp between (**E**) *prospero*-*doublesex* and (**F**) *prospero*-*UXT*. (**G**) Strength of selection on *doublesex* OD1 domain estimated from branch-site models in codeml (dN/dS < 1, blue; dN/dS = 1, black; dN/dS > 1, orange) and Godon (LRT test *p* < 0.05 after BH correction indicated by red circles on edges).

**Figure 4:**
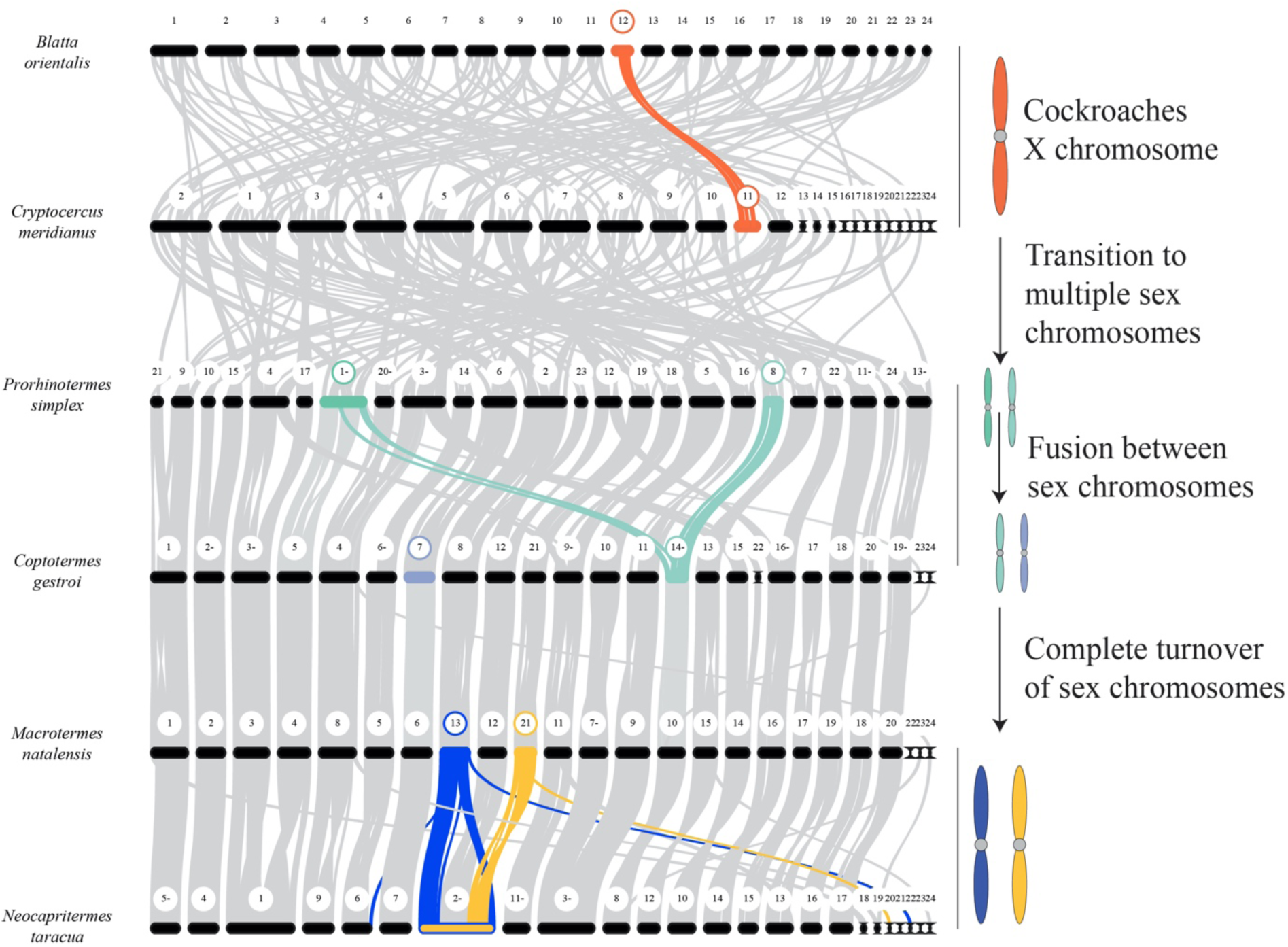
Turnover of sex chromosomes in termites. The turnover of identity was assessed through macrosyntenic gene patterns among the longest scaffolds of cockroaches and selected Neoisoptera. The X chromosome of cockroaches became autosomal in termites. Sex chromosome identity shows frequent turnover among termite lineages and was fixed in Termitidae. In *N. taracua*, the two arms occur on the same chromosome (see Figure 2 **I**) suggesting either a fusion or a mis-assembly considering the conserved Geoisopteran karyotype comprising *n* = 21 chromosomes.

We hypothesized that the translocation of *dsx* in Mastotermitidae, Archotermopsidae, and Hodotermopsidae, initiated the formation of a Y chromosome. To test this hypothesis, we determined whether *dsx* resides on sex-linked scaffolds or autosomes by using our scaffold classification (Supplementary Data 4,10) and by mapping male and female reads to species-specific *dsx* sequences (Figure 3C; Supplementary Data 12). Our results indicated that *dsx* is autosomal in the two cockroaches and in most termite genomes (Figure 3; Supplementary Data 11,12). The RNA splicer *tra* and *tra-*2 and other transcription factors regulating the sex-determining cascade in other insects, such as *Sex-lethal* (*Sxl*) and *virilizer* (*vir*) (*24*, *48*), are also autosomal in most Icoisoptera (Supplementary Data 13). In contrast, *dsx* was only found in males in Mastotermitidae, Archotermopsidae, Hodotermopsidae, and Rhinotermitidae (Figure 3; Supplementary Data 11,12). In Mastotermitidae, *dsx* was located on scaffold #51, a small 860-kbp scaffold not part of the 40 scaffolds making up the L90, and for which 96% of mapping contigs from the male-only short-read assembly exhibited male coverage (log_2_(*C_MF_*) > 0.85; Dryad File 2). Scaffold #51 likely represents a fragment of the Y chromosome. Finally, while *dsx* was missing from the assemblies of *H. sjostedti* and *Z. nevadensis*, it was found in the raw nanopore reads, indicating that *dsx* was dropped during the co-assembly of the minimally divergent X and Y chromosomes (Supplementary Data 4). Altogether, our results suggest that the presence of *dsx* on an autosome-looking sex chromosome, rather than chromosome-wide differences, determines sex through male-restricted expression in early-diverging termite lineages, raising the possibility that it was the condition in the ancestor of termites. Overall, the X and Y chromosomes of Mastotermitidae, Hodotermopsidae, and Archotermopsidae are undifferentiated and fully recombine, as evidenced by karyotype-wide presence of chiasmata between homologous chromosome pairs (Figure 1H). The inheritance of *dsx*, which acts as the initiator of the sex-determining cascade, determine male development in these termites (Figure 5).

**Figure 5:**
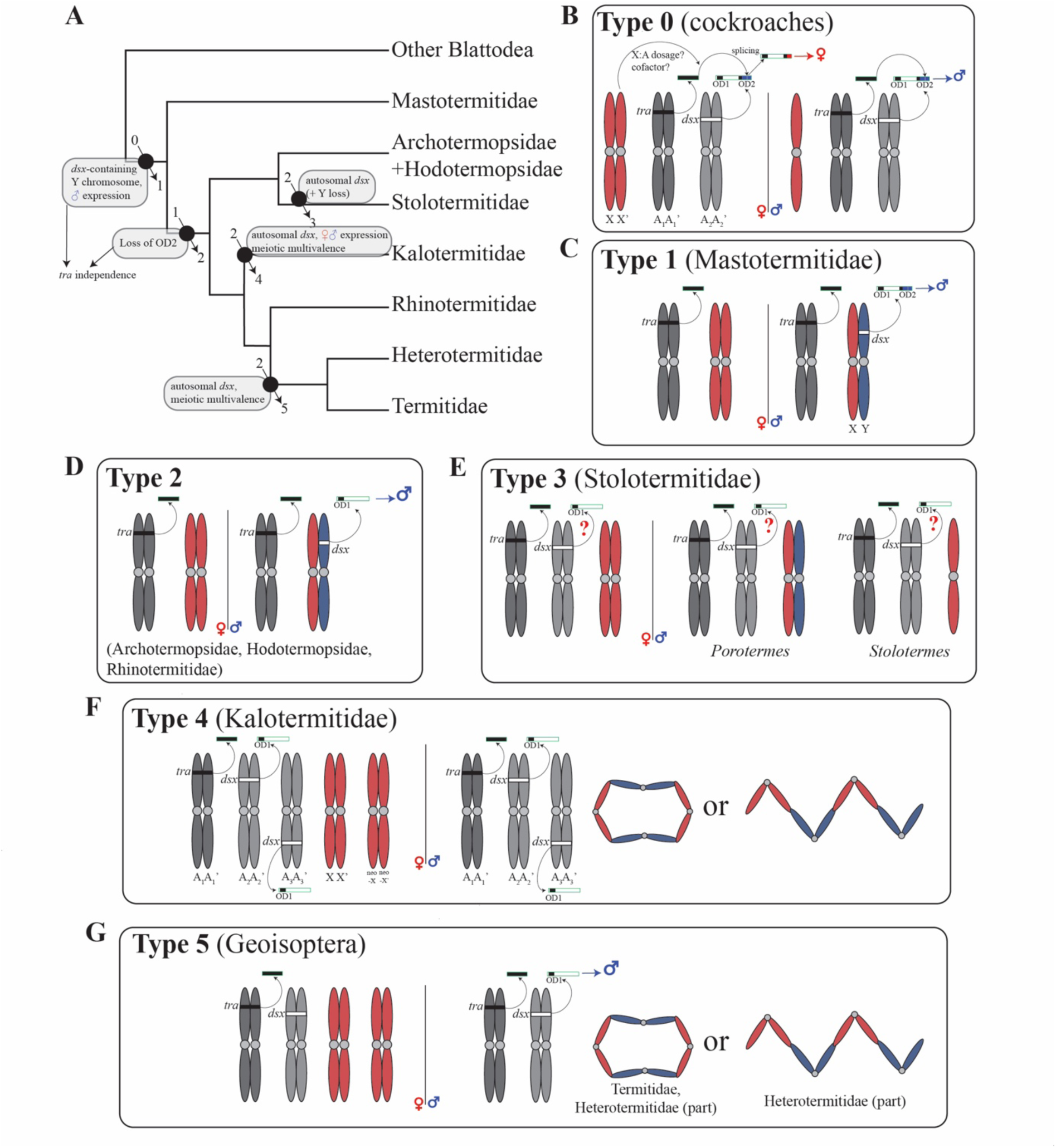
Major transitions of termite sex determination systems. We classify sex determination systems (SDS) among six broad types (numbered 0-5) according to the SDS and genomic location of sex-determining genes (*doublesex*, *dsx*; *transformer*, *tra*). (**A**) Simplified phylogenetic relationships among families studied herein (adapted from the UCE-based classification tree of Hellemans *et al*. (*120*) mapping the possible transitions among the main SDS types. (**B**) Type 0 (non-termite Blattodea): males are hemizygous for the X chromosome, and the female sex is determined by *tra* through splicing of *dsx*, while male is determined by full-length *dsx*. Whether splicing is influenced by the X chromosome-to-autosome ratio remains unknown. Termites (SDS types 1-5) evolved an XX/XY SDS. Sex is determined by *dsx* located on the Y chromosome in (**C**; Type 1) Mastotermitidae and (**D**; Type 2) Archotermopsidae, Hodotermopsidae and Rhinotermitidae. In other types, *dsx* is autosomal and the SDS remains vastly uncharacterized. The expression patterns remain unknown in Stolotermitidae (**E**; Type 3), expressed by both sexes in Kalotermitidae (**F**; Type 4), and in males in Geoisoptera (**G**; Type 5). Type 4 and 5 present the occurrence of chains and rings of chromosomes during meiosis in males. The loss of the *dsx* OD2 domain at the transition between type 1 and other types indicates that sex determination may not rely on *tra* splicing, although *tra* may still be required for mRNA maturation of *dsx*.

The existence of a single undifferentiated sex chromosome in early-diverging termite lineages does not necessarily mean a recent origin (Figure 1). Some organisms have retained ancient homomorphic sex chromosomes. For instance, Z and W chromosomes are largely homologous in ratite birds and sturgeon fishes despite being 130 and 180 My old, respectively (*49*, *50*). These old homomorphs are possibly maintained by weak sexual selection, limiting divergence and sexual antagonistic polymorphisms (*1*). Considering that sexual selection is weak or nonexistent in termites (*51*), it is not excluded that the sex chromosomes of Mastotermitidae, Archotermopsidae, and Hodotermopsidae have an ancient origin. Future studies examining the homologies between the sex chromosomes of these lineages may help resolve the origin of these homomorphs.

### Sex chromosomes experienced frequent turnover across non-Termitidae termites

Our genomic analyses revealed the diversity of termite SDSs. Termites present lineage-specific variations in the location of *dsx* (on autosomes or the Y chromosome), the presence of XX/XY and XX/X0 SDSs, and the presence of one or multiple sex chromosomes interacting during male meiosis (Figures 2-3). These observations suggest turnover of sex chromosomes amongst termite lineages. We assessed sex chromosome turnover in termites by examining their homology through shared macrosynteny blocks on the near chromosome-scale assemblies of the two cockroaches and 11 Neoisoptera with identified sex chromosomes (Figure 4; Supplementary Figures 5-6). Our analyses indicated that the identity of sex chromosomes is variable between families of Neoisoptera, within non-Termitidae Neoisoptera families, and sometimes even within species. Firstly, the sex chromosomes of *C. gestroi* (Heterotermitidae) do not share large-scale homology with the sex chromosomes of *M. natalensis* and other Termitidae (Figure 4), nor do they carry evident sex-determining genes (Supplementary Data 14). This first suggest that the exact sex-determining mechanisms likely differ across termite families. Secondly, only one of the two sex chromosomes of *C. gestroi* is homologous with one of the two sex chromosomes of *Prorhinotermes simplex* (Psammotermitidae). Thirdly, the identity and number of sex chromosomes vary within Heterotermitidae, as exemplified by the absence of homology between *C. gestroi* and *R. flavipes* sex chromosomes (Supplementary Figure 6; Dryad: File 2), with *R. flavipes* possessing more sex-linked scaffolds (scaffolds 8, 14, 18, and 20; Dryad: File 2; Supplementary Figures 3,4). These results are consistent with the variable meiotic figures found in *Reticulitermes*, varying between bivalents and octavalents in *Reticulitermes lucifugus* (Figure 1 L,M) (*29*, *52*). Finally, none of the sex chromosomes we identified in termites match the X chromosome that *B. orientalis* and *Cryptocercus meridianus* share (Figure 4), confirming a complete turnover of sex chromosomes in the ancestor of termites, as previously speculated from short-read assemblies (*53*). The lack of homology between the sex chromosomes of termites and their sister groups extends to other parts of the genomes, as all chromosomes experienced extensive genomic rearrangements as the lineages branching first in the termite tree diverged from their common ancestor (Supplementary Figure 6). Therefore, termite genomes, including sex chromosomes, exhibit structural variability, the extent of which remains to be fully characterized.

### Sex chromosome identity is conserved across multiple subfamilies of Termitidae

We investigated chromosome homology through gene synteny blocks between *M. natalensis* and six species of Termitidae with chromosome-level genome assemblies. Our results indicated that the two sex chromosomes of *M. natalensis* (scaffolds 13 and 21; Figure 2J, K) are homologous to, and syntenic across their whole length with, the two sex chromosomes of *Odontotermes formosanus*, *Promirotermes redundans*, and *Neocapritermes taracua*, in which the two sex chromosomes appear fused with one autosome (Supplementary Figure 5; Dryad: File 2). In two other species, *Anoplotermes pacificus* and *Coatitermes* sp., we were able to identify only one clear sex chromosome among the L90 scaffolds (Supplementary Data 1; Dryad: File 2). The scaffold 21 of *M. natalensis* was homologous to the identified sex chromosome of *A. pacificus* and to two smaller scaffolds in the assembly of *Coatitermes* (scaffolds 21 and 22, the latter being sex-linked; Supplementary Figure 5; Dryad: File 2). Finally, *Sphaerotermes sphaerothorax* did not exhibit any major sex-linked scaffold (Dryad: File 2), possibly due to shallower sequencing depth, but two scaffolds matched the two sex chromosomes of *M. natalensis* (Supplementary Data 1). Because two sex chromosomes were expected in all Termitidae (Supplementary Data 2), it is likely that the differentiation of some sex chromosomes was below our detection threshold. Therefore, our results indicated that the two sex chromosomes retained homology across at least five Termitidae subfamilies, with some potential variations, such as the fusion of the two sex chromosomes and one autosome in *N. taracua*, although this may be a mis-assembly. Our results suggested that Termitidae have largely retained the same sex chromosomes for 50 My, since they diverged from their last common ancestor. The conserved sex chromosomes of Termitidae contrast with the karyotypic variability of sex chromosomes among species of Kalotermitidae (*29*, *30*, *35*, *54*).

### Sex chromosomes and dsx may have acquired additional functions linked to termite sociality

The turnover of sex chromosomes is well-documented in various organisms possessing a single sex chromosome, such as frogs and fishes (*16*, *55*). Our results show that most termite species have also undergone sex chromosome turnover, but a turnover that involves multiple sex chromosomes, often comprising a large fraction of the genome, as seen in *Glyptotermes nakajimai*, in which 90% of the genome is sex-linked (*56*). Termite sex chromosome turnover was proposed to as a mechanism that increases whole genomic diversity, potentially providing a selective advantage to organisms that often have an inbred lifecycle (*56*, *57*). Following this hypothesis, the turnover of sex chromosomes allows the heterozygosity accumulated by males on the sex chromosomes (*58*, *59*) (Figures 1G, 2E) to become autosomal. The turnover of sex chromosomes would thus act as a “fountain of youth,” releasing male-accumulated heterozygosity across both sexes. Additional studies are needed to test this model.

Besides initiating male development, *dsx* has been proposed to have evolved additional functions, including in the caste developmental pathway, which would explain the presence of several *dsx* paralogs with bisexual expression in some termites, such as the Kalotermitidae (*41*). We investigated the evolution of the *dsx* OD1 domain using dN/dS analyses and attempted to identify the genes it regulates. Our codeml analysis revealed high dN/dS values for branches leading to the Icoisoptera and the families of Neoisoptera, suggesting ancient episodes of positive or relaxed selection (Figure 3G). Our Godon analysis further revealed a significant dN/dS increase in Termitidae, a family that retained homologous sex chromosomes (Figure 4), suggesting that some genes present in the SSRs could be differentially regulated by *dsx* between sexes. We attempted to identify the genes regulated by *dsx* in *M. natalensis* by locating specific DNA-binding motifs using HOMER. Our analyses identified 859 genes possibly regulated by *dsx*, including 25 located in the SSRs (Supplementary Data 15). GO term analyses of these 25 genes indicated links to spermatogenesis (2 genes), oogenesis (5 genes), and DNA-binding (11 genes) (Supplementary Data 16). Notable examples include the Ecdysone-induced 74EF transcription factor, which is expressed during post-meiotic stages of spermatogenesis (*60*), and the *sarah* protein, which is critical for meiotic progression in oocytes (*61*, *62*). Our results support the role of *dsx* in spermatogenesis- and longevity-related processes, as previously hypothesized (*40*). More importantly, 22 of the 25 genes were linked to cellular processes (GO:0009987), including four with cell cycle functions (GO:0007049): the mitotic (GO:0000278) and meiotic (GO:0051321) cell cycles, the centrosome cycle (GO:0007098), and the anchoring of microtubules at the centrosome (GO:0034454). These results suggest the involvement of *dsx* in the regulation of genes involved in the sex-specific segregation of sex chromosomes during meiosis (Figure 2).

Finally, the *dsx* transcription factor may also be associated with caste differentiation mechanisms in termites. A major change undergone by *dsx* was the loss of its sex-specific OD2 domain in all termite species surveyed in this study and in previous studies (*38–41*), except in *Mastotermes* (Supplementary Data 17), in which OD2 is expressed only in males (accessions XXX). The male-specific expression of OD2-containing *dsx* transcripts is unique to *Mastotermes*, differentiating it from a sex determination based on the sex-specific splicing of OD2 as in cockroaches, and its loss as in other termites (Figure 3,5). Notably, all ants except the basal Ponerinae also lost the OD2 domain (*63*), suggesting potential convergent mechanisms between ants and termites in their caste determination mechanisms. While domain gains and rearrangements have been found to be associated with the evolution of eusociality, domain losses have not been studied appropriately (*64*). The loss of the sex-specific domain OD2 is predicted to lead to reduced DNA-binding specificity (*65*, *66*), which in turn leads to less differentiated males and females. Here, we posit that *dsx* may have been co-opted for caste-determining mechanisms in termites, which could explain the frequent caste-sex specialization in Termitidae (*67–71*). This hypothesis is in line with *dsx* regulating juvenile hormone signaling in the stag beetle (*72*), an hormone central to insect embryogenesis and termite caste determination (*73–75*). Further studies are required to clarify the role of *dsx* in termite caste development, for example, through controlled RNAi experiments.

### Conclusions

Our results show that sex chromosomes present a low level of differentiation, which likely reflects ongoing recombination between largely homologous sex chromosomes (Figures 1, 2) and frequent sex chromosome turnover (Figure 4). Therefore, homology and gene content are largely retained between X and Y chromosomes, suggesting minimal dosage compensation, at least in non-Termitidae termites, contrasting previous reports (*53*). Phased assemblies of X and Y chromosomes will be required to further resolve these patterns. In early-diverging termites, sex is determined by the presence of *dsx* on the Y chromosome, while *dsx* is mostly dissociated from the sex chromosomes in Icoisoptera, indicating modification of the *tra*-*dsx* sex determination pathway (Figures 3,5). Our results further suggest that *dsx* may be involved in the formation of novel sex-related genomic structures during meiosis, thereby facilitating the accumulation of heterozygosity in termite males. Transcriptomics combined with RNAi experiments will be required to clarify the role of *dsx* during termite development.

## Material and Methods

### 1. Biological samples and sequencing

We used the genome assemblies produced by Liu *et al*. (*42*) for the identification of termite sex chromosomes. We focused on a subset of 36 species (25 of which were scaffolded assemblies, including 21 near chromosome-level), for which pools of males and females were separately sequenced (Supplementary Data 1). One male and one female adult were sequenced for both species of cockroaches. We sequenced male and female reproductives and reproduction-destined individuals (*i.e.*, nymphs, alates, primaries, or neotenics) for 23 termite species, male and female worker-derived individuals (*i.e.*, workers, soldiers) for 10 termite species, and a queen and a pool of male and female workers for the termitid species *Anoplotermes pacificus* (for details, see Supplementary Data 1). The sex of individuals was determined based on the configuration of sternites with a stereomicroscope.

The paired-end 150-bp reads were either reused from Liu *et al*. (*42*) or newly generated in this study (for details, see Supplementary Data 3). For reused reads, the library preparation was either carried out using the NEBNext Ultra II FS DNA kit or the Tell-Seq WGS library prep kit, and sequencing was performed on the Illumina HiSeq X or NovaSeq 6000 platforms, respectively. For newly generated paired-end 150-bp reads, libraries were prepared using the NEBNext Ultra II FS DNA library prep kit with one-fifth of the manufacturer’s recommended reagent volumes. Libraries were sequenced on the Illumina HiSeq X platform. Newly generated raw reads were deposited on the NCBI Sequence Read Archive (SRA) under the BioProject accession number XXXXX. A complete list of the reads used in this study is available in Supplementary Data 3. For all downstream analyses, raw reads were trimmed to remove adapters and low-quality bases using fastp *v*0.20.1 (*76*).

### 2. Identification of sex chromosomes

We aimed to identify sex chromosomes for each of the 36 species for which sex-specific short reads were available (Supplementary Data 1). Because termite sex chromosomes appear poorly differentiated on karyotypes (*31*), we used a total of six computational methods relying on comparative male-female coverage (three methods), SNPs (two methods), and k-mers (one method). This integrative approach minimizes the number of false positives.

#### 2.1. Identification of sex chromosomes from comparative coverage

We used three differential coverage approaches to identify a set of candidate sex-linked scaffolds. For each species, we mapped sex-specific trimmed reads onto (*i*) the repeat-masked reference genome assemblies Liu *et al*. (*42*) (reference-based approach; *e.g.*, (*77*)); (*ii*) male genome assemblies generated from male short reads (male-genome approach; *e.g.*, (*7*)); and (*iii*) the longest transcripts of all genes annotated in the reference genome assemblies (transcript-based approach; *e.g.*, (*78*)). For method (*ii*), we generated a male genome assembly using male trimmed reads. Trimmed reads were normalized to an average depth of 50X and removing reads with a below apparent depth of 5 using the BBNorm tool from BBMap *v*38.86 (*79*), and assembled using metaSPAdes *v*3.15.1 (*80*). For all methods, male and female trimmed reads were separately mapped using BWA-MEM *v*0.7.10 (*81*) and sorted BAM alignment files were generated using the ‘fixmate’ and ‘sort’ commands of SAMtools *v*1.9 (*82*).

Male-to-female coverage ratios (*C_MF_*) allow to differentiate autosomal chromosomes and pseudo-autosomal regions of the X and Y chromosomes from X- and Y-specific regions of the sex chromosomes (*7*, *55*). We used the normalized male and female coverage from the repeat-masked reference genome assemblies (method *i*), male genome assemblies (method *ii*), and the longest transcripts of all genes annotated in the reference genome assemblies (method *iii*) to estimate *C_MF_*. For the analyses performed on reference genomes and male genome assemblies (methods *i* and *ii*), the normalized scaffold coverage was obtained from BAM files using the ‘idxstats’ and ‘flagstat’ commands of SAMtools. The ‘idxstats’ command provided the number of reads mapping each scaffold, and the ‘flagstat’ command provided the total number of reads mapping all scaffolds of the genome assembly. Normalized scaffold coverage was obtained by dividing the number of reads mapping to a given scaffold by the total number of reads mapping to the entire genome assembly. For the transcript-based method (*iii*), we calculated *C_MF_* using the male and female normalized mean transcript coverage. The mean transcript coverage was obtained using the ‘coverage’ command of BEDtools *v*2.29.2 (*83*) with the arguments “-mean -sorted”. The mean transcript coverage was then divided by the mode coverage of all transcripts for normalization (*78*). Transcripts with a coverage three-fold higher than the mode coverage were filtered out.

The normalized male coverage was divided by the normalized female coverage to obtain *C_MF_* values for each scaffold (methods *i* and *ii*) and transcript (method *iii*). Autosomal and pseudo-autosomal regions are expected to exhibit equal depth of coverage in both sexes (*C_MF_ ≈* 1; log_2_(*C_MF_*) *≈* 0). In contrast, when the X and Y chromosomes are fully differentiated, the depth of coverage of the X chromosome is expected to be twice as high in females as in males (*C_MF_ ≈* 0.5; log_2_(*C_MF_*) *≈ -*1), while the coverage of Y-specific regions is expected to be zero in females (*C_MF_ =* ∞; log_2_(*C_MF_*) *=* ∞). The X and Y chromosomes can exhibit little or no differentiation from one another. When sex chromosomes present no differentiation, their coverage is similar to that of autosomes (*C_MF_ ≈* 1; log_2_(*C_MF_*) *≈* 0), while their coverage falls between that of autosomes and fully differentiated sex chromosomes under partial differentiation, leading to intermediate *C_MF_* values. Here, we considered scaffolds or transcripts with log_2_(*C_MF_*) values falling in the interval ]-∞,-0.65] as putatively X-linked, ]-0.65,0.85[ as putatively autosomal or pseudo-autosomal, and [0.85,+∞[ as putatively Y-specific. Note that this approach, based on comparative male-female coverage, may fail to identify sex chromosome scaffolds and transcripts in case of highly reduced or lack of differentiation between X and Y chromosomes. Scaffolds and transcripts considered autosomal and pseudo-autosomal by these methods may be part of undifferentiated sex chromosomes, whose identification requires additional approaches.

The method *i* directly used scaffolds from the reference genome assembly of Liu *et al*. (*42*) and directly flagged them as sex chromosome scaffolds or not. In contrast, the methods *ii* and *iii* flag scaffolds from the male genome assembly (method *ii*) or transcripts (method *iii*) as part of the sex chromosomes or the autosomes. One additional step is required to link these scaffolds and transcripts to the scaffolds of the reference genomes. Scaffolds from the male genome assemblies with a length of at least 5 kbp were matched to the reference genome using megaBLASTn v2.13.0+ (*84*) with the arguments “-max_target_seqs 1 -max_hsps 1”. We considered reference scaffolds to be part of the sex chromosomes when at least 10% of the scaffolds mapping to them were sex-linked (male or female). Transcripts were ported to their corresponding scaffold using the reference genome annotation GFF3 file. We only considered reference scaffolds containing at least 10 genes. Reference scaffolds of termite genomes were considered part of the sex chromosomes when at least 2% of their associated transcripts were sex-linked. We applied this low threshold for termite genomes due to the homomorphic nature of their sex chromosomes (*31*), suggesting that they are largely undifferentiated. Reference scaffolds of cockroach genomes (*Blatta orientalis* and *Cryptocercus meridianus*) were considered part of the sex chromosomes when at least 50% of their associated transcripts were sex-linked. This higher threshold was appropriate for cockroaches and their XX/X0 SDS.

#### 2.2. Identification of sex chromosomes from SNPs

Comparative coverage approaches can fail to identify sex chromosomes when X and Y chromosomes exhibit low levels of divergence, in which case they co-assemble. One alternative approach to identify such undifferentiated sex chromosomes is the use of SNPs. Heterogametic males are expected to exhibit a higher number of SNPs than females on their sex chromosomes, as regions that recently stopped recombining on the Y chromosomes accumulate SNPs independently from the X chromosomes (*85*). Therefore, SNPs-based methods can help identifying weakly divergent sex chromosomes exhibiting *C_MF_* values similar to autosomes (*i.e.*, nascent sex chromosomes). The method *iv* identified sex-linked scaffolds with higher number of SNPs in heterogametic males compared to females (SNPs-density approach; *e.g.*, (*77*)). The method *v* identified sex-linked scaffolds based on their content in biallelic SNPs homozygous in females and heterozygous in males, which is the SNP heterozygosity pattern expected on the sex chromosomes of an XX/XY SDS (SNPs-coherence approach; *e.g.*, (*86*)).

We performed variant calling on the reference genome assemblies of Liu *et al*. (*42*) with either male or female BAM alignment files using the ‘mpileup’ command of bcftools *v*1.9 (*87*). To associate allelic frequencies to variant calls, we ran ‘mpileup’ using the flag ‘--annotate FORMAT/AD,FORMAT/ADF,FORMAT/ADR,FORMAT/DP,FORMAT/DP4,FORMAT/SP ,INFO/AD,INFO/ADF,INFO/ADR’ followed by ‘bcftools +fill-tags’. This procedure creates variant BCF files containing all additional fields available. The variant BCF files were normalized using ‘bcftools norm’ with the flags ‘--fasta-ref,’ which left-align and normalize indels, and ‘-m -any,’ which split multiallelic variants into biallelic variants. The normalized BCF files were then filtered with “bcftools view -i ’%QUAL>=20’ -e ’(FORMAT/DP4[:0]+FORMAT/DP4[:1]+FORMAT/DP4[:2]+FORMAT/DP4[:3])<=10’” to remove variants with quality scores < 20 and read depths < 10, and “bcftools filter --SnpGap 10” to filter out SNPs located within 10-bp of indels. SNPs were extracted from the resulting files using the command “bcftools view --types snps” (hereafter: the SNPs BCF file). Finally, we used “bcftools view --max-alleles 2” to only retain biallelic SNPs and “bcftools view -i ’(FORMAT/AD[:0])/(FORMAT/AD[:0]+FORMAT/AD[:1]) >= 0.3 && (FORMAT/AD[:0])/(FORMAT/AD[:0]+FORMAT/AD[:1]) <= 0.7’” to only retain variants with allelic depth falling between 0.30 and 0.70. The last step ensures that retained SNPs are true heterozygous sites and removes artifacts resulting from the uneven sequencing of pools of males or females (hereafter: the biallelic SNPs BCF file).

The SNPs-density approach (method *iv*) relied on the estimation of a male-to-female SNP ratio (*SNP_MF_*), which is expected to be higher for sex chromosomes than autosomes. To calculate *SNP_MF_*, we counted the number of raw SNPs per scaffold in males and females separately using the biallelic SNPs BCF files. The per-scaffold SNP count in males and females was divided by their respective genome-wide sequencing depths for depth-normalization. The male-to-female SNP ratio (*SNP_MF_*) was calculated by dividing male normalized SNP counts by female normalized SNP counts. We only performed this analysis for scaffolds exhibiting at least 10 SNPs across both sexes and 1 SNP in males. Scaffolds were considered sex-linked when they exhibited at least twice more SNPs in males than females (log_2_(*SNP_MF_*) ≥ 1).

The SNPs-coherence approach (method *v*) attempts to identify sex chromosome scaffolds based on the proportion of SNPs matching an XX/XY SDS, that is SNPs that are homozygous in females and heterozygous in males. We focused on SNP positions identified in the biallelic SNPs BCF file and recorded their homozygosity/heterozygosity status in males and females in four files. For each sex, we used “bcftools view” with the flags (a) “--genotype hom” on the SNPs BCF file to extract all homozygous genomic positions (homozygous BCF file), and (b) “--genotype het” on the biallelic SNPs BCF file to obtain all SNPs with two alleles (heterozygous BCF file). We used the male heterozygous BCF file and the female homozygous BCF file to obtain the species coherence BCF file that contained all SNPs biallelic in males and homozygous in females, *i.e.*, composed only of the genomic positions compatible with an XX/XY SDS (hereafter: coherent SNPs). We obtained the coherent SNPs through four successive steps. First, we extracted the list of genomic positions contained in the male heterozygous BCF file using “bcftools query -f ’%CHROM\t%POS\n”. Second, we used the obtained list of positions in males to extract the corresponding positions from the female homozygous BCF file with the command “bcftools view”. Third, we combined the male and female SNPs into one line per genomic position using “bcftools merge”. This intermediary file contains heterozygous sites in males, as well as sites that are homozygous or with unknown status (*i.e.*, either non-homozygous or not meeting the quality thresholds) in females. The fourth and last step created the species coherence BCF file by filtering out all genomic positions with unknown status in females with the command “bcftools view -e ’GT[*] = "mis"’”, and therefore only contains the coherent SNPs. We used this species coherence BCF file to count the number of coherent SNPs per scaffold, considering only scaffolds with 10 or more coherent SNPs. We obtained the coherent-to-male SNPs ratio (*SNP_XY_*) for a given scaffold by dividing the number of coherent SNPs by the total number of SNPs recorded in the biallelic SNPs BCF files of both males and females. In this SNPs-coherence approach (method *v*), we defined scaffolds exhibiting at least 50% of coherent SNPs (*SNP_XY_* ≥ 0.50) as sex chromosomes. For cockroaches, we considered scaffolds to be sex linked when *SNP_XY_* ≤ 0.10.

Termites have a XX/XY SDS. However, for the sake of completeness, we also performed the SNPs-coherence validation approach (method *v*) for species following a ZW/ZZ. We obtained the number of incoherent SNPs per scaffold, that is, the number of SNPs compatible with the species following a ZW/ZZ (female heterozygous/male homozygous) SDS, using the above-outlined approach but with the female heterozygous and male homozygous BCF files.

#### 2.3. Identification of sex chromosomes from Y-mers

The method *vi* was a k-mer counting method that searches for male-specific k-mers (*i.e.*, Y-mers) across read sets of both sexes (k-mer approach; *e.g.*, (*7*, *88*)). Trimmed paired-end reads sets (R1 and R2 files) were merged using the BBMerge tool from BBMap separately for males and females (*89*). To avoid counting undefined k-mers, merged reads containing undefined “N” nucleotides were removed using SeqKit *v*0.13.2 (*90*).

Several operations were carried out to identify Y-mers with sufficient coverage. First, we identified all Y-mers sequences using unikmer *v*0.19.1-1+b3. Canonical k-mers were counted separately in males and females using a k-mer length of 31 bp with the ‘count’ command of unikmer. K-mers shared by males and females were identified using the command ‘sort --repeated’ of unikmer. Sex-restricted k-mers were identified by comparing sex-specific k-mers and shared k-mers using the ‘diff’ command of unikmer, and fasta sequences were extracted with the ‘view’ command (hereafter: the unimers files). Second, we filtered out low-coverage k-mers using jellyfish *v*2.2.7 (*91*). We extracted canonical 31-bp k-mer sequences from male reads using the ‘count’ and ‘dump’ commands of jellyfish and filtered out k-mers with a coverage below 30 (hereafter: the coverage files). Third, we identified unique Y-mers with coverage > 30 by comparing the unimers and coverage files using the commands ‘count’, ‘sort --repeated’ and ‘view’ of unikmer (hereafter: the Y-mer files). Fourth, we extracted the merged fasta reads containing unique Y-mers. We used the ‘uniqs’ command of unikmer with the flags ‘-M -m 150,’ which identifies merged reads of at least 150 bp that can contain multiple Y-mers. We extracted the corresponding merged fastq reads using seqtk *v*1.3 (https://github.com/lh3/seqtk). Fifth, we mapped the extracted merged fastq reads to the reference genome using BWA-MEM and the ‘sort’ command of SAMtools and counted the number of Y-mers-containing reads (hereafter: the Y-mer reads) for each scaffold using the ‘idxstats’ command of SAMtools. Finally, we calculated the Y-mer ratio (*R_Y_*) of each scaffold, which was the number of Y-mer reads on this scaffold divided by the mean number of Y-mer reads of all scaffolds of that genome assembly. We considered termite scaffolds as being male-linked when *R_Y_* ≥ 5. For cockroaches, we considered scaffolds to be sex-linked when *R_Y_* ≤ 5.

#### 2.4. Consensus sex chromosome set

We defined a consensus set of sex chromosomes by summarizing the results from the six computational methods described above. A scaffold was considered sex-linked when most methods identified it as sex-linked. A scaffold with an equal number of methods considering it sex-linked and autosomal was not considered sex-linked. We used the classification of scaffolds, as sex-linked or autosomal, for all downstream analyses.

### 3. dN/dS analyses of autosomes and sex chromosomes

Genes present on sex chromosomes may experience different selection regimes than autosomal genes, reflecting the total or partial suppression of recombination in sex chromosomes or sex-specific selection (*43–45*). We used non-synonymous to synonymous substitution rates (dN/dS) to investigate the strength of selection across autosomes and sex chromosomes. We compared the patterns of dN/dS values of genes present on sex chromosomes to those of genes present on autosomes.

We used the 34,594 hierarchical orthogroups (HOGs) identified with OrthoFinder *v*2.5.2 (*92*) by Liu *et al*. (*42*) to calculate dN/dS ratios. We used the longest isoforms found in the 47 genomes. We retained HOGs containing at least four genes and fewer than two genes per species on average. This approach removed the very small and large HOGs, leaving a total of 17,652 HOGs. Protein sequences were aligned using MAFFT *v*7.505 (*93*) with all parameters left under default settings. Protein alignments were used to build phylogenetic trees using IQ-TREE *v*2.0.7 (*94*) run with the optimal substitution model selected by ModelFinder (*95*). Long branches, which often represent outliers, were detected and removed from the trees and protein alignments using TreeShrink *v*1.3.9 (*96*) with a quantile cutoff of 0.01. Protein alignments were converted into codon alignments with PAL2NAL *v*14 (*97*). Codon alignments were used to reconstruct final HOG trees with IQ-TREE using the optimal codon substitution model identified by ModelFinder. dN/dS ratios were calculated for each branch of the gene trees using the codeml free-ratio model (model = 1; NSsites = 0) from PAML *v*4.9 (*98*). We restricted our analysis to the branches of the trees having dS values between 0.01 and 10, as branches with extreme values can lead to unreliable estimations of dN/dS ratios.

We compared gene dN/dS values between consensus autosomal and sex-linked scaffolds using two-sided Student *t*-tests implemented in the ‘t.test’ function of the R package ‘stats.’ We restricted our analyses to species having at least 20 sex-linked genes. We performed two-sided Student *t*-tests on 28 species and corrected the *p*-values for multiple comparisons using the Benjamini-Hochberg false discovery rate (“method = BH”) with the ‘p.adjust’ function from the R package ‘stats.’

### 4. Evolutionary strata along and among sex chromosomes

We identified sex chromosomes using sex-specific reads for 22 species with chromosome-level genome assemblies published by Liu *et al*. (*42*) (Supplementary Data 1). For these sex chromosomes, we aimed to distinguish between sex-specific regions (SSRs) —which display accumulated sequence divergence between males and females owing to recombination suppression—, and the pseudo-autosomal regions (PAR), which still undergo recombination. We only considered chromosomes that made up the L90 set.

We determined *C_MF_* values and the number of coherent SNPs along 5-kbp and 50-kbp windows of each chromosome composing the L90 set, respectively. We defined windows for each reference genome independently using the ‘makewindows’ command of BEDtools. The *C_MF_* values of 5-kbp windows were calculated using the male and female sliding window coverage, each obtained with the ‘coverage’ command of BEDtools run with the arguments “-mean -sorted” on the BAM alignment and window files. The number of coherent SNPs along 50-kbp windows was obtained using the ‘intersect’ command of BEDtools on the species coherence BCF and window files.

### 5. Syntenic map of sex-linked chromosomes across species

The synteny among chromosome-level genome assemblies was visualized using the MCscan module of JCVI v1.4.19 (*99*) run on the genome annotations of Liu *et al*. (*42*). More specifically, GFF3 genome annotation files were converted into BED files and mRNA features were extracted using the “jcvi.formats.gff” module of JCVI. Pairwise alignments were performed using the longest transcript of each gene with the lastal command (*100*) of the last-align v1447-1+b1 package of Debian-Med (*101*). The syntenic map was visualized using the “jcvi.graphics.karyotype” module of JCVI with the argument “--dist=20” for a default chaining distance of 20.

### 6. Chromosome imaging

We prepared termite meiotic karyotypes following the protocol of Luykx (1990). Male gonads were dissected in 1% sodium citrate and incubated therein for 10 minutes. Gonads were then transferred onto a microscope slide and spread in a first fixative, a glacial solution of acetic acid:ethanol:water (1:1:1.3). Prior to the full desiccation of tissues, a few drops of a second fixative, a glacial acetic acid:ethanol (1:1) solution, were added, and slides were immediately transferred into a Coplin jar containing the final fixative, a glacial acetic acid:ethanol (1:3) solution, in which they were incubated for at least 20 min. Slides were dehydrated using an ethanol series of 75%, 95%, and 100% ethanol. The slides were then stained with a 1:60 Giemsa’s Azur-Eosin-Methylene Blue (AppliChem) solution in Gurr buffer (Gibco) for 8 min. Slides were visualized on a Nikon Eclipse Ni microscope with a 40x dry objective. Images were acquired with a mounted Nikon DS-Fi1c camera.

### 7. The tra-dsx sex determination cascade

#### 7.1. Identifying doublesex

The *doublesex* (*dsx*) gene encodes a transcription factor with a DNA-binding DM (OD1) domain common to *Doublesex* and *Mab-3* Related Transcription (DMRT) genes, and a sex-specific dimerization (OD2) domain (*65*, *102*, *103*). We conducted tblastN searches across all 47 genomes to identify the OD1 and OD2 domains of the *dsx* transcription factor. We carried out tblastN searches on the entire genome assemblies to identify the OD1-containing sequences (DMRT genes and *dsx*) using known *dsx* and DMRT protein sequences as queries (Supplementary Data 7). The OD2 domain was then searched using tblastN with a protein sequence of OD2 (PFAM: PF08828) extracted from the *dsx* gene of *Cryptocercus punctulatus* (accession BCX65399). Resulting matches were extracted with BEDtools *v*2.29.2 (*83*).

We further characterized *dsx* and identified the features distinguishing it from other DMRT factors by searching the genomes with the extracted OD1-containing nucleotide sequences using HMMER3 v3.1b2 (http://hmmer.org; (*104*)). Our tblastN searches of OD1 indicated the presence of *dsx* and four other DMRT genes (DMRT11, DMRT93, DMRT99, and an uncharacterized DMRT-related gene, DMRTX; Supplementary Data 10). Sequences were translated into all six possible amino acid frames using the “translate6frames.sh” tool of BBMap, and the OD1 domain HMM profile (accession PF00751) from the Pfam database (*105*, *106*) was used as a query for the HMMER searches. HMMER searches indicated that the OD1 domain was absent from the DMRTX gene. Our tblastN-HMMER searches did not return any match to the OD1 domain of *dsx* in two genomes; however, it was found in the raw nanopore reads using minimap2 *v*2.20 (*107*) with the arguments “-ax map-ont” and their species-specific sequences as query (*H. sjostedti*: LC635719; *Z. nevadensis*: BR002455). Extracted nanopore reads were assembled with Flye *v*2.9.5 (*108*).

We reconstructed a maximum-likelihood (ML) tree of the OD1 domain to distinguish *dsx* from other DMRT (*i.e.*, DMRT11, DMRT93, and DMRT99) genes (Supplementary Figure 7). We used reference sequences for each gene (Supplementary Data 8), and the best-hit frame of each sequence from the HMMER search. Sequences were aligned using MAFFT *v*7.305, and the tree was reconstructed with IQ-TREE *v*2.2.2.5 (*94*) under default settings.

We investigated the synteny of the *dsx*, sandwiched between the homeobox *prospero*-like (*pros*; containing the PF05044 homeo-prospero domain; CDD: 461534) and the prefoldin *UXT* (UXT; PFAM: PF02996; CDD: cd23158) genes in termites (*20*, *38*, *41*). We reannotated these genes because some were not annotated in the consensus annotations of Liu *et al*. (*42*). First, we identified the scaffold on which each gene was located. For paralogs of the *dsx* gene, we used the genome coordinates of the OD1 domain as identified in the tree-based approach to differentiate it from other DMRT genes. For the other two genes, we used tblastN on each of the 47 genomes using the amino acid sequences of PSN43307 (*prospero*) and PSN43305 (*UXT*) from *Blatta germanica* as queries. Second, we retrieved 3–13 nucleotide sequences (Supplementary Data 9), translated them into protein sequences with the “transeq” function of EMBOSS *v*6.6.0 (*109*), and constructed each protein HMM profile using HMMER3. Finally, we used the HMM profiles to reannotate the three genes on the extracted scaffolds with the “genome mode” of bitacora v1.4 (*110*) that enables GeMoMa v1.7 (*111*, *112*). The reannotations of these genes are provided on the Dryad repository (File 5).

#### 7.2. Rates of evolution of the *dsx* gene

We investigated the strength and direction of selection operating on the OD1 domain of *dsx* during termite evolution. We used the OD1 sequences identified in the 47 reference genomes of Liu *et al*. (*42*), together with published OD1 sequences of *H. sjostedti* (GenBank accession: LC635719) and *Z. nevadensis* (BR002455), two species for which OD1 sequences could not be retrieved from the reference genomes. DNA sequences were translated into protein sequences using the “transeq” command of EMBOSS and aligned with MAFFT. The protein alignments were converted into codon alignment with PAL2NAL. We reconstructed a ML tree using the codon alignment of the OD1 domain and IQ-TREE run with a GTR+G substitution model. Gene tree-species tree reconciliation was performed using the species tree of Liu *et al*. (*42*) and GeneRax v2.1.3 (*113*) run with a GTR+G model of nucleotide substitution. To investigate signatures of positive selection, we analyzed each branch of the reconciled tree using the codeml program implemented in PAML v 4.9 (*98*) and Godon v2021-09-13 (*114*). For the codeml analysis, we used the free-ratio model (model = 1) to directly estimate one dN/dS ratio per branch (*115*). Elevated dN/dS ratios were interpreted as indicative of positive selection. For the Godon analysis, branch lengths were first estimated with the codon alignment and a M0 model (*116*), which assumes a single dN/dS ratio across all alignment sites and tree branches.

These branch length estimates served as the null hypothesis and were compared to dN/dS ratios estimated with a branch-site model incorporating codon-level gamma rates, the BSG model (*117*). Likelihood-ratio tests were performed to assess whether the BSG model provided a significantly better fit than the M0 model for each focal branch. A significantly better fit of the BSG model was interpreted as evidence for positive selection. To control for multiple comparisons, we calculated the false discovery rate (FDR) using the Benjamini-Hochberg method to correct p-values.

#### 7.3. Genes regulated by *dsx*

We used the scaffolded genome of *Macrotermes natalensis* as a reference to attempt to identify genes whose transcription is regulated by *dsx*. We did so by locating specific DNA-binding motifs throughout the genome using HOMER v5.1 (*118*). We used the nucleotide sequences previously used to find these genes in *Reticulitermes speratus* (*40*). We converted each DNA-binding motif FASTA sequence into a motif HOMER file with the “seq2profile.pl” tool and allowed one mismatch to account for potential positive selection on the binding domain OD1 at the basis of Termitidae. We combined all motifs into a single motif file with which we screened the genome using the “scanMotifGenomeWide.pl” tool. We only considered DNA-binding hits found less than 3-kbp upstream of each gene feature of interest recorded in the genome GFF3 annotation file. We intersected DNA-binding hits to all upstream regions using BEDtools (Supplementary Data 15).

Genes putatively regulated by *dsx* were functionally annotated based on homology to proteins from the RefSeq genomes of *Drosophila melanogaster* (GCF_000001215.4), *Periplaneta americana* (GCF_040183065.1), *Zootermopsis nevadensis* (GCF_000696155.1), and *Cryptotermes secundus* (GCF_002891405.2). The protein FAA and annotation GFF3 files of each genome were retrieved using EDirect v18.2. Each candidate gene was searched against each genome’s protein sequences separately with BLASTX implemented in BLAST 2.13.0+ (*84*). Gene Ontology (GO) terms were obtained by querying protein accession with the ‘query’ function of the ‘mygene’ R package (*119*). GO terms were further classified into biological processes (BP) or molecular functions (MF) using the ‘GO.db’ R package with the ‘GOBPOFFSPRING’ and ‘GOMFOFFSPRING’ functions, respectively (Supplementary Data 16). BP terms included cellular processes (GO:0009987), cell cycle (GO:0007049), microtubule cytoskeleton organization (GO:0000226), spermatogenesis (GO:0007283), and oogenesis (GO:0048477). MP terms contained nucleic acid binding (GO:0003676) and DNA-binding transcription factor activity (GO:0003700).

#### 7.4. Genes activating *dsx*

We screened the 47 genomes published by Liu *et al*. (*42*) for the sequences involved in the *transformer*-*dsx* sex determination cascade. We conducted tblastN searches to identify (*i*) the *transformer* (*tra*) gene (Accession: QGB21094; *Blattella germanica*), (ii) *tra-2* (KDR12437; *Z. nevadensis*), (*iii*) the sex-lethal *Sxl* gene (XP_021921630; *Z. nevadensis*), and (*iv*) the *virilizer* mRNA cofactor (XP_021932582; *Z. nevadensis*).

We searched for homologous regions in the shared sex-linked scaffolds of *Coptotermes gestroi* and *M. natalensis* in order to identify potential candidates involved in *dsx* male-specific transcription pattern in Geoisoptera (Supplementary Data 14). We reannotated these regions *de novo* using a bidirectional best BLAST hit approach implemented in BLAST 2.13.0+ (*84*) in order to avoid missing any protein of interest and because some gene features were missing from the consensus annotation of Liu *et al*. (*42*) (*e.g.*, the *dsx* gene was absent in a few genome assemblies while present in the reads). Homologous regions were identified using megablastN with the options “-max_hsps 1 -max_target_seqs 1”. Protein-coding sequences were identified with tblastN using reference proteins from the four above-mentioned RefSeq genomes. Our candidate proteins were annotated using BLASTX matches with e-value ≤ 1e-5 and coverage and identity ≥ 30, as described above. Only hits of at least 100bp in megablastN, tblastN, and BLASTX searches were retained.

## Acknowledgments

We thank the Sequencing Section (SQC) and the Scientific Computing & Data Analysis Section (SCDA) of OIST for assistance with sequencing and providing access to the OIST computing cluster, respectively. This work was supported by subsidiary funding from OIST. SH was supported by the Japan Society for the Promotion of Science (JSPS) through a postdoctoral fellowship for foreign researchers (19F19819) and as a Postdoctoral Researcher of the Fonds de la Recherche Scientifique – FNRS. JŠ was supported by the project IGA No. 20253134 from the Faculty of Tropical AgriSciences of the Czech University of Life Sciences Prague. AB was supported by the Czech Science Foundation (GAČR) grant Junior STAR No. 23-08010M.

## Author Contributions Statement

SH conceptualized the experiments. EK performed lab experiments and generated data. SH, YMW, and KF analyzed the data. MJ karyotyped termites. SH and TB wrote the original draft manuscript. All authors edited and accepted the final version of this manuscript.

## Data availability

Samples used in this study are described in Supplementary Data 1. The raw sequence reads generated in this study have been deposited in the NCBI Sequence Read Archive under the BioProject accession code PRJNAXXXXXXX [https://www.ncbi.nlm.nih.gov/bioproject/ PRJNAXXXXXXX]. The raw data from prior publications used in this study are available under BioProject accession codes PRJNAXXXXXXX [https://www.ncbi.nlm.nih.gov/bioproject/ PRJNAXXXXXXX], PRJNAXXXXXXX [https://www.ncbi.nlm.nih.gov/bioproject/ PRJNAXXXXXXX], PRJNAXXXXXXX [https://www.ncbi.nlm.nih.gov/bioproject/ PRJNAXXXXXXX]. The raw sequence data can also be accessed using the individual accession numbers given in Supplementary Data 3. Additional data is also available in the Dryad Digital Repository at: XXXX.

## Code availability

No new custom code was published with this article.

